# The *Aedes aegypti* peritrophic matrix controls arbovirus vector competence through HPx1, a heme–induced peroxidase

**DOI:** 10.1101/2022.06.02.494599

**Authors:** Octavio A. C. Talyuli, Jose Henrique M. Oliveira, Vanessa Bottino-Rojas, Gilbert O. Silveira, Patricia H. Alvarenga, Asher M. Kantor, Gabriela O. Paiva-Silva, Carolina Barillas-Mury, Pedro L. Oliveira

## Abstract

*Aedes aegypti* mosquitoes are the main vectors of arboviruses. The peritrophic matrix (PM) is an extracellular layer that surrounds the blood bolus and acts as an immune barrier that prevents direct contact of bacteria with midgut epithelial cells during blood digestion. Here, we describe a heme-dependent peroxidase, hereafter referred to as heme peroxidase 1 (HPx1). HPx1 promotes PM assembly and antioxidant ability, modulating vector competence. Mechanistically, the heme presence in a blood meal induces HPx1 transcriptional activation mediated by the E75 transcription factor. HPx1 knockdown increases midgut reactive oxygen species (ROS) production by the DUOX NADPH oxidase. Elevated ROS levels reduce microbiota growth while enhancing epithelial mitosis, a response to tissue damage. However, simultaneous HPx1 and DUOX silencing was not able to rescue bacterial population growth, as explained by increased expression of antimicrobial peptides (AMPs), which occurred only after double knockdown. This result revealed hierarchical activation of ROS and AMPs to control microbiota. HPx1 knockdown produced a 100-fold decrease in Zika and Dengue 2 midgut infection, demonstrating the essential role of the mosquito PM in the modulation of arbovirus vector competence. Our data show that the PM connects blood digestion to midgut immunological sensing of the microbiota and viral infections.

## Introduction

Mosquito-borne viruses are emerging as global threats to public health. Female mosquitos ingest infected blood from a host and transmit the virus to another host during the next blood-feeding. As the first insect tissue infected by the virus, the midgut is the initial barrier that the virus must overcome to establish itself in the mosquito (Black IV et al., 2002). Because blood digestion occurs in the midgut concomitantly with viral infection of epithelial cells, digestion-triggered physiological events have a major influence on the course of intestinal infection (Talyuli et al., 2021).

The peritrophic matrix (PM) in mosquitoes is a semi-permeable chitinous acellular layer secreted by intestinal cells after blood feeding. The PM completely envelopes the blood bolus, and its structure avoids direct contact of the digestive bolus with the midgut epithelia (Lehane, 1997; Shao et al., 2001). The PM is the site of deposition of most of the heme produced from blood hemoglobin hydrolysis, thus limiting exposure of the midgut cells to harmful concentrations of heme, a pro-oxidant molecule (Pascoa et al., 2002). Extensive gut microbiota proliferation occurs in most hematophagous insects after a blood meal. Therefore, the PM is a barrier that limits interaction of the tissue with the intestinal microbiota (Kuraishi et al., 2013; Oliveira et al., 2011; Terra et al., 2018), playing a role analogous to the mammalian intestinal mucous layer (Terra et al., 2018). The PM is mainly composed of chitin and proteins, and correct assembly of this structure is crucial to its barrier function. Additionally, the PM is a barrier for parasites such as *Plasmodium, Trypanosoma brucei*, and *Leishmania major*, which must attach to or traverse the PM to complete their development in an insect vector (Coutinho-Abreu et al., 2010; Rose et al., 2020; Shahabuddin et al., 1995; Weiss et al., 2014).

There are several studies on the role of reactive oxygen species (ROS) and redox metabolism on the gut immune response to pathogens. In *Drosophila melanogaster*, ROS production by a dual oxidase enzyme (DUOX, an NADPH oxidase family member) is triggered by pathogenic bacteria. The self-inflicted oxidative damage arising from DUOX activation is prevented by hydrogen peroxide scavenging via an immune-regulated catalase (IRC) (Ha, Oh, Bae, et al., 2005; Ha, Oh, Ryu, et al., 2005). In *Anopheles gambiae, Plasmodium* ookinete midgut invasion triggers a complex epithelial response mediated by nitric oxide and hydrogen peroxide that is crucial to mount an effective mosquito antiplasmodial response (Kumar & Barillas-Mury, 2005). Furthermore, an *Anopheles gambiae* strain genetically selected to be refractory to *Plasmodium* infection exhibits enhanced activation of JNK-mediated oxidative stress responses (Garver et al., 2013; Jaramillo-Gutierrez et al., 2010). In *Aedes aegypti*, it has been proposed that the Dengue NS1 viral protein decreases hydrogen peroxide levels, preventing an oxidative intestinal environment, which is an adverse condition for both Dengue and Zika viral infection (Bottino-Rojas et al., 2018; Liu et al., 2016). Catalase silencing in the *Aedes aegypti* gut reduces the dengue infection prevalence rate (Oliveira et al., 2017). The ROS generation by DUOX plays a key role in modulating proliferation of the indigenous microbiota, growth of opportunistic pathogenic bacteria, and Dengue virus infection (Ha et al., 2009; Liu et al., 2016; Oliveira et al., 2011).

Kumar et al. (2010) showed that heme peroxidase 15 (HPx15), also referred to as immunomodulatory peroxidase (IMPer), is expressed in the *A. gambiae* midgut and uses the hydrogen peroxide generated by DUOX as a substrate to crosslink proteins of the mucous layer in the ectoperitrophic space, limiting diffusion of immune elicitors from the gut microbiota and thus preventing activation of midgut antimicrobial responses to commensal bacteria. IMPer silencing results in constant activation of epithelial immune responses against both bacteria and *Plasmodium* parasites (Kumar et al., 2010). A similar immune barrier role for the PM against parasite infection has also been shown in tsetse flies infected with *T. brucei* and sandflies infected with Leishmania (Ramalho-Ortigao, 2010; Rose et al., 2020). Therefore, most of the studies on the PM of insect disease vectors have focused on its role as a barrier for parasites, but much less is known about the influence of PM on viral infections or its contribution to gut homeostasis and immune responses in *A. aegypti*.

Here, we show that HPx1, a heme peroxidase associated with the *A. aegypti* PM, has a dual role, acting in the PM assembly crucial for its barrier function and as an antioxidant hydrogen peroxide-detoxifying enzyme. This role of HPx1 in midgut physiology and immunity highlights that dietary heme is a signal that by triggering HPx1 expression and PM function, produces a homeostatic response that controls ROS and AMP immune effectors, microbiota expansion, and viral infection.

## Materials and Methods

### Ethics Statement

All the animal care and experimental protocols were conducted following the guidelines of the institutional care and use committee (Comissão de Ética no Uso de Animais, CEUA-UFRJ) and the NIH Guide for the Care and Use of Laboratory Animals. The protocols were approved under the registry #CEUA-UFRJ 149/19 for rabbit use and 075/18 for mice immunization and antiserum production. The animal facility technicians at the Instituto de Bioquímica Médica Leopoldo de Meis (UFRJ) carried out all aspects related to rabbit and mice husbandry under strict guidelines to ensure humane animal handling.

### Mosquitos

The *Aedes aegypti* females (Red-Eye strain) used in this study were raised in an insectary of Universidade Federal do Rio de Janeiro. Approximately 200 larvae were reared in water-containing trays and fed dog chow. Pupae were transferred to plastic cages, and adults were fed *ad libitum* with 10% sucrose solution in cotton pads. The insects were kept in a 12 h dark/light period-controlled room at 28 °C and 80% humidity. Blood feeding was performed using rabbit ears or artificially through glass feeders sealed with Parafilm and connected in a circulated water bath at 37 °C. Substitute of blood meal (SBM) is a previously described artificial diet with a chemically defined composition (Talyuli et al., 2015) and was used in experiments in which the presence of heme was modulated. For this study, only bovine albumin and gamma-globulin were used as protein sources, and no hemoglobin was added. Hemin was solubilized in 0.1 M NaOH and neutralized with 0.01 M sodium phosphate buffer (pH 7.4). Antibiotics (penicillin 200 U/ml and streptomycin 200 μg/mL) in autoclaved 5% sucrose solution were supplied for 3 days before feeding with blood or SBM.

### Catalase activity

Midguts were dissected in cold 50% ethanol, and epithelia were separated from the blood bolus surrounded by the PM. The samples were immediately transferred to tubes with a protease inhibitor cocktail (50 μg/mL SBTI, 1 mM benzamidine, 1 mM PMSF). The midgut epithelial samples were directly homogenized, but the PM-enriched fraction samples were centrifuged 3x at 10000 × g for 5 min at 4 °C to remove as much of the blood bolus as possible. Hydrogen peroxide detoxification activity was measured based on peroxide absorbance (240 nm for 1 min) in the presence of mosquito homogenates (Aebi, 1984), and the protein concentration was determined according to Lowry (LOWRY et al., 1951). For *in vitro* inhibition experiments, samples were incubated with different concentrations of 3-amino-1,2,4-triazole for 30 min at 4 °C before enzymatic activity assays (Oliveira et al., 2017).

### Double-strand RNA synthesis and injections

To synthesize dsRNA, a first PCR was performed using mosquito whole-body cDNA as a template. The product was diluted 100x and used in a second reaction with T7 primers. dsLacZ was used as a control and amplified from a cloned plasmid containing the LacZ gene. Double-strand RNA synthesis was performed using a MEGAscript T7 transcription kit (Ambion/Thermo Fisher Scientific, MA - USA). The reaction was performed overnight at 37 °C. Each product was precipitated with 1 volume of isopropanol and 1:10 volume of 3 M sodium acetate (pH 3). Four-day-old females were cold-anesthetized, and a double shot of 69 nl of 3 μg/μL dsRNA was injected into the mosquito thorax using a Nanoject II (Drummond Scientific, PA - USA). For double-silencing experiments, the dsRNA mixture containing both dsHPx1 and dsDUOX was lyophilized and then resuspended to half of the original volume. One day after injection, the females were blood-fed.

### RNA extraction, cDNA synthesis, and qPCR

Total RNA was extracted from midgut samples using TRIzol reagent following the manufacturer’s protocol. The RNA (1 μg) was treated for 30 min at 37 °C with DNAse, and cDNA was synthesized with a High-Capacity cDNA Reverse Transcription Kit (Applied Biosystems/Thermo Fisher Scientific, MA - USA) using random primers. Quantitative PCR (qPCR) was performed with a StepOnePlus Real-Time PCR System (Applied Biosystems/Thermo Fisher Scientific, MA - USA) using Power SYBR-green PCR master MIX (Applied Biosystems/Thermo Fisher Scientific, MA - USA). The RP49 gene was used as an endogenous control, and the primers used in the qPCR analyses are listed in Supplemental Table 1.

### HPx1 antiserum

*Aedes aegypti* HPx1 (AAEL006014) was cloned into pET15b at the NdeI (5’) and BamHI (3’) restriction sites. The export signal predicted by SignalP software (Petersen et al., 2011) was removed from the purchased codon-optimized sequence (GenScript, NJ - USA). Plasmid pET15b containing the HPx1 protein-coding sequence was transformed into the *Escherichia coli* BL21(DE3) strain. Cells were grown at 37 °C in 2xYT medium containing 100 μg/L ampicillin. After reaching OD_600_ 0.4, the cells were cooled and supplemented with 0.5 mM isopropyl β-D-1-thiogalactopyranoside (IPTG), and the cells were incubated at 25 °C overnight. The cells were then harvested and resuspended in buffer A (200 mM Tris pH 8.0, 500 mM NaCl, 5 mM imidazole, 1% Triton X-100, 10% glycerol, 10 mM β-mercaptoethanol) supplemented with 2 mg/mL lysozyme. After breaking the cells by ultrasonic treatment, the insoluble fraction was collected by centrifugation. Because the recombinant HPx1 obtained by this protocol was not soluble, the corresponding protein band was cut from the SDS–PAGE gel, and the protein was extracted from the gel (Retamal et al., 1999). Immunization of BalB/C mice was performed by injecting two shots of 50 and 25 μg of antigen intraperitoneally, spaced by 21 days, using Freund’s complete and incomplete adjuvant, respectively, in a 1:1 ratio (1 antigen:1 adjuvant). Two weeks after the second shot, blood was extracted by cardiac puncture, and the serum was isolated and frozen for further use.

### HPx1 Western Blotting

Pools of midgut epithelia and PMs were dissected and immediately placed in tubes containing a protease inhibitor cocktail. The samples were denatured at 95 °C in the presence of SDS sample buffer, and a volume equivalent to 1 midgut/slot was resolved by SDS–PAGE. The gel was blotted onto PVDF membranes (Bjerrum Schafer-Nielsen buffer - 48 mM Tris, 39 mM glycine, 0.037% SDS - pH 9.2, 20% methanol) for 1 h at 100 V and blocked with 5% albumin in TBS-T (Tris 50 mM, pH 7.2, NaCl 150 mM, 0.1% Tween 20) overnight (ON) at 4 °C. The membranes were incubated with 1:5000 anti-HPx1 primary antiserum diluted in blocking solution for 5 h at room temperature. The primary antibody solution was removed, and the membrane was washed with TBS-T (3X) before incubation with alkaline phosphatase-conjugated anti-mouse secondary antibody (1:7500 in blocking solution) for 1 h at room temperature (RT). The membrane was washed and developed using NBT/BCIP alkaline phosphatase substrates.

### Putative HPx1 gene promoter *in silico* analysis

The MEME motif-based algorithm (Bailey et al., 2009) was used to analyze a 2500 bp sequence upstream of the 5’ transcription start site of the HPx1 gene (*Aedes aegypti* genome, version AaegL3.4). The Fimo tool (Grant et al., 2011) was used to specifically search for cis-regulatory elements associated with ecdysone molecular signaling, as previously described (Kokoza et al., 2001).

### PM permeability

HPx1-silenced mosquitoes were artificially fed rabbit blood containing 1 mg/ml dextran-FITC (Sigma, FD4), and the midguts were dissected 18 h after feeding. The midguts were fixed in 4% paraformaldehyde solution in 0.1 M cacodylate buffer for 2 h and placed ON in a 15% sucrose solution in phosphate-buffered saline (PBS; 10 mM Na-phosphate buffer, pH 7.2, 0.15 M NaCl) at RT. Then, they were incubated in a 30% sucrose solution for 30 h. The following day, the midguts were infiltrated with 50% Optimal Cutting Temperature/OCT (Tissue-TEK, Sakura Finetek, CA - USA) in 30% sucrose solution for 24 h and then ON in 100% OCT. The samples were frozen at -70°C until use, and serial 14-μm-thick transverse sections were obtained using an MEV Floor Standing Cryostat (SLEE Medical, Nieder-Olm - Germany). The slices were placed on glass slides and mounted with Vectashield containing DAPI mounting medium (Vector laboratories, CA - USA). The sections were examined in a Zeiss Observer.Z1 with a Zeiss Axio Cam MrM Zeiss through excitation BP 546/12 nm, beam splitter FT 580 nm, and emission LP 590 nm.

### Reactive oxygen species measurement

Midguts were dissected at 18 h after blood feeding and incubated with 50 μM dihydroethidium (hydroethidine; DHE; Invitrogen/Thermo Fisher Scientific, MA - USA) diluted in 5% fetal bovine serum-supplemented Leibovitz 15 medium for 20 min at room temperature in the dark. The dye medium was removed, and the midguts were gently washed in dye-free medium. The fluorescence of the oxidized DHE was acquired using a Zeiss Observer.Z1 with a Zeiss Axio Cam MrM Zeiss, and the data were analyzed using AxioVision software in a Zeiss-15 filter set (excitation BP 546/12 nm; beam splitter FT 580 nm; emission LP 590 nm) (Oliveira et al., 2011).

### Mitosis labeling

HPx1-silenced mosquitoes were blood-fed, and the midguts were dissected 18 h after feeding. The midguts were fixed in 4% paraformaldehyde solution for 30 min, permeabilized with 0.1% Triton X-100 for 15 min at RT, and blocked ON at room temperature in a solution containing PBS, 0.1% Tween 20, 2.5% BSA, and 10% normal goat serum. All samples were incubated overnight with a mouse anti-PH3 primary antibody (1:500, Merck Millipore, Darmstadt - Germany) diluted in blocking solution at 4 °C and then washed 3x for 20 min each in washing solution (PBS, 0.1% Tween 20, 0.25% BSA). The midguts were incubated with a secondary goat anti-mouse antibody conjugated with Alexa Fluor 546 (Thermo Fisher Scientific, MA - USA) for 1 h at room temperature at a dilution of 1:2000, and nucleic acids were stained with DAPI (1 mg/ml, Sigma) diluted 1:1000. PH3-positive cells were visualized and counted using the Zeiss fluorescence microscope mentioned above (Taracena et al., 2018).

### Virus infection and titration

Zika virus (ZKV; Gen Bank KX197192) was propagated in the *Aedes albopictus* C6/36 cell line for 7 days in Leibovitz-15 medium (Gibco #41300–039, Thermo Fisher Scientific, MA - USA; pH 7.4) supplemented with 5% fetal bovine serum, tryptose 2.9 g/L, 10 mL of 7.5% sodium bicarbonate/L; 10 mL of 2% L-glutamine/L, 1% nonessential amino acids (Gibco #11140050, Thermo Fisher Scientific, MA - USA) and 1% penicillin/streptomycin (Oliveira et al., 2017) at 30 °C. Dengue 2 virus (DENV, New Guinea C strain) was propagated in C6/36 in MEM media (GIBCO #11095080, Thermo Fisher Scientific, MA - USA) supplemented with 10% fetal bovine serum and 1% penicillin/streptomycin for 6 days (Jupatanakul et al., 2017). The cell supernatants were collected, centrifuged at 2,500 × g for 5 min, and stored at -70 °C until use. Mosquitoes were infected with 10^5^ PFU/ml ZKV or 2×10^7^ PFU/ml DENV in a reconstituted blood meal prepared using 45% red blood cells, 45% of each virus supernatant, and 10% rabbit serum (preheated at 55 °C for 45 min). Four days after Zika infection or seven days after Dengue infection, the midguts were dissected and stored at -70 °C in 1.5 ml polypropylene tubes containing glass beads and DMEM supplemented with 10% fetal bovine serum and 1% penicillin/streptomycin. The samples were thawed and homogenized, serially diluted in DMEM, and incubated in 24-well plates with a semiconfluent culture of Vero cells (for ZKV samples) or BHK21 cells (for DENV) for 1 h at 37 °C and then incubated with DMEM 2% fetal bovine serum + 0.8% methylcellulose (Sigma, M0512) overlay for 4 days at 37 °C and 5% CO_2_ in an incubator. The plates were fixed and stained for 45 min with 1% crystal violet in ethanol/acetone 1:1 (v:v).

### Statistical analyses and experimental design

All experiments were performed in at least three biological replicates, and samples correspond to pools of 5–10 insects. The graphs and statistical analyses were performed using GraphPad 8 software. For qPCR experiments, relative gene expression was calculated by the Comparative Ct Method (Pfaffl, 2001), and the result is expressed as the mean of ΔΔCt values compared to a housekeeping gene (RP-49, AAEL003396) (Gentile et al., 2005).

## Results

### *Aedes aegypti* PM detoxifies hydrogen peroxide

After a blood meal, the antioxidant capacity of the *Aedes aegypti* midgut is increased by expressing enzymes and low molecular weight radical scavengers (Sterkel et al., 2017). These protective mechanisms are complemented by the capacity of the PM to sequester most of the heme produced during blood digestion, which has been proposed to be a preventive antioxidant defense, as heme is a pro-oxidant molecule (Devenport et al., 2006; Pascoa et al., 2002). Figure 1A shows that *A. aegypti* PM exhibited hydrogen peroxide detoxifying activity up to 24 h ABM followed by a sharp decrease at 36 h, close to the end of blood digestion. The specific activity of the PM hydrogen peroxide scavenging activity was comparable to the activity found in the midgut epithelia 24 h after blood-feeding, which is attributed to a canonical intracellular catalase (SUP1A). However, silencing of the “canonical” intracellular catalase (AAEL013407-RB) did not alter the PM’s ability to detoxify hydrogen peroxide at 24 h after feeding (Fig 1B), suggesting that this activity in the PM is not due to the midgut intracellular catalase (AAEL013407-RB) (Oliveira et al., 2017). This hypothesis received support from the observation that the hydrogen peroxide decomposing activity of the PM and midgut epithelia showed distinct *in vitro* sensitivity to the classical catalase inhibitor amino triazole (SUP1B). Moreover, neither depletion of the native microbiota by antibiotic treatment (SUP1C) nor feeding the insect an artificial diet (cell-free meal) devoid of catalase (SUP1D) altered PM hydrogen peroxide detoxification, additionally excluding the hypothesis of an enzyme originating from the microbiota or host red blood cells.

**Figure 1.**
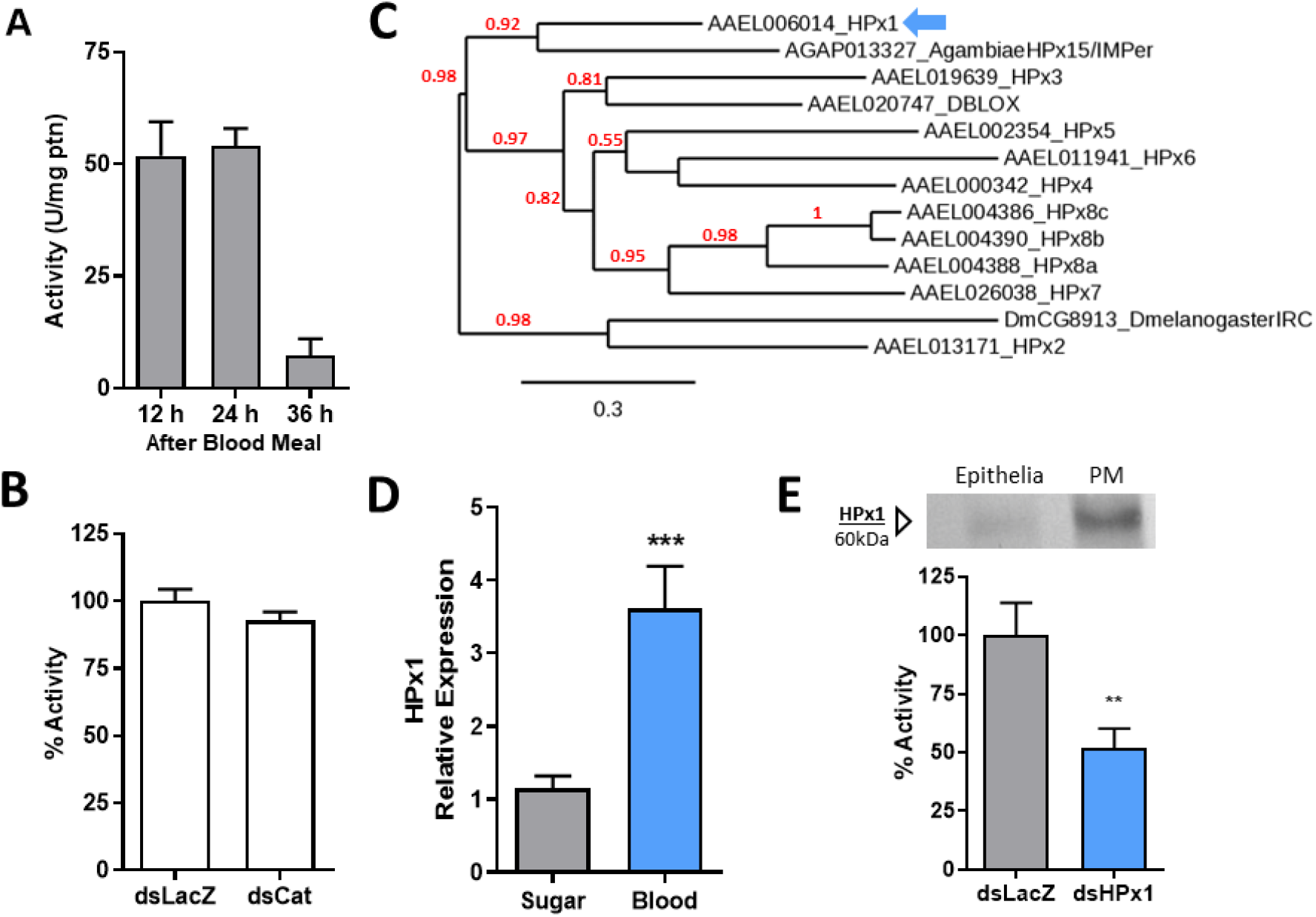
Hydrogen peroxide scavenging by the *Aedes aegypti* peritrophic matrix. A) PMs were dissected at different times after a blood meal (ABM), and their catalase specific activity was measured (12 h n=3, 24 h n=3; 36 h n=2). B) Catalase activity of PMs dissected from control (dsLacZ-injected) and catalase (AAEL013407-RB)-knockdown insects at 24 h AMB (dsLacZ n=9; dsCat n=9). C) Phylogenetic tree of heme peroxidases from *Aedes aegypti* (AAEL), *Anopheles gambiae*, and *Drosophila melanogaster*. Maximum likelihood analysis was performed, and the numbers in each branch represent bootstraps. (D) HPx1 expression in midguts at 24 h ABM relative to the sugar-fed control (Sugar n=11; Blood n=11). (E) Western blot of the HPx1 protein in 20 μg of gut epithelia and PM extracts at 18 h ABM and catalase activity of PMs dissected from control and HPx1-knockdown insects (dsLacZ n=8; dsHPx1 n=13). The full western blot membrane is shown in Supplementary Figure 1E. Data are the mean +/-SEM. **p> 0.005, ***p<0.001 for the T test for D and E.

### HPx1-dependent hydrogen peroxide scavenging by the PM

We hypothesized that the observed PM hydrogen peroxide detoxifying activity should be attributed to another enzyme encoded by the mosquito genome. Peroxidases are a multigene family of enzymes and the genome of *A. aegypti*, as with most other organisms, has many peroxidases. Peroxidases are grouped in three large families: glutathione, heme, and thioredoxin peroxidases. As the PM is an extracellular structure, we initially searched for peroxidases with a predicted signal peptide. Interestingly, this search identified ten peroxidases, all of them belonging to the heme peroxidase family, which also includes the secreted peroxidases of *Anopheles gambiae* (Kumar et al., 2010) and *Drosophila melanogaster* (Ha, Oh, Ryu, et al., 2005). Phylogenetic analysis showed that *A. aegypti* heme peroxidase 1 (HPx1) is a close homolog of the peroxidase HPx15/IMPer that promotes the crosslink of extracellular proteins in the gut lumen of *A. gambiae* (FIG 1C). As the HPx15/IMPer from *A. gambiae* was shown to be secreted by the midgut epithelia, we used the presence of a secretion signal peptide as an additional feature to indicate HPx1 as the *A. aegypti* PM enzyme responsible for decomposing hydrogen peroxide. Fig. 1D shows that the blood meal induced HPx1 gene expression in the gut. Moreover, western blotting showed that most of the HPx1 in the midgut was bound to the PM, with a minor fraction being associated with the epithelia (Fig. 1 E), and RNAi silencing of HPx1 expression significantly decreased hydrogen peroxide detoxification by the PM (Fig. 1 E; HPx1 silencing efficiency: SUP1F).

### The E75 transcription factor mediates heme-induced HPx1 midgut expression

A blood meal triggers large changes in the gene expression pattern of *A. aegypti* and, among several factors, the heme released upon hemoglobin proteolysis acts as a pleiotropic modulator of transcription (Bottino-Rojas et al., 2015). Feeding the insects with SBM with or without heme revealed that heme significantly regulated HPx1 gene expression (Fig. 2A). Accordingly, lower hydrogen peroxide decomposition activity was observed in the PM secreted by females fed SBM without heme compared to the blood-fed insects, a phenotype rescued by heme supplementation (Fig. 2B). *In silico* analysis of the HPx1 promoter gene region revealed putative binding sites for E75, a hormone-responsive transcription factor that functions as a heme and redox sensor (SUP2A) (Cruz et al., 2012). E75 knockdown significantly reduced HPx1 gene expression in the midgut after blood feeding, revealing a molecular mechanism for triggering HPx1 by heme (Fig. 2C; E75 silencing efficiency: SUP2B). Proliferation of the gut microbiota in response to blood feeding is known to induce expression of several genes in the midgut. However, neither microbiota depletion by oral administration of antibiotics (aseptic) nor reintroduction of a bacterial species (*Enterobacter cloacae*) into antibiotic-treated mosquitoes affected HPx1 gene expression (SUP2C). Thus, we propose the molecular signaling model shown in Fig. 2D.

**Figure 2.**
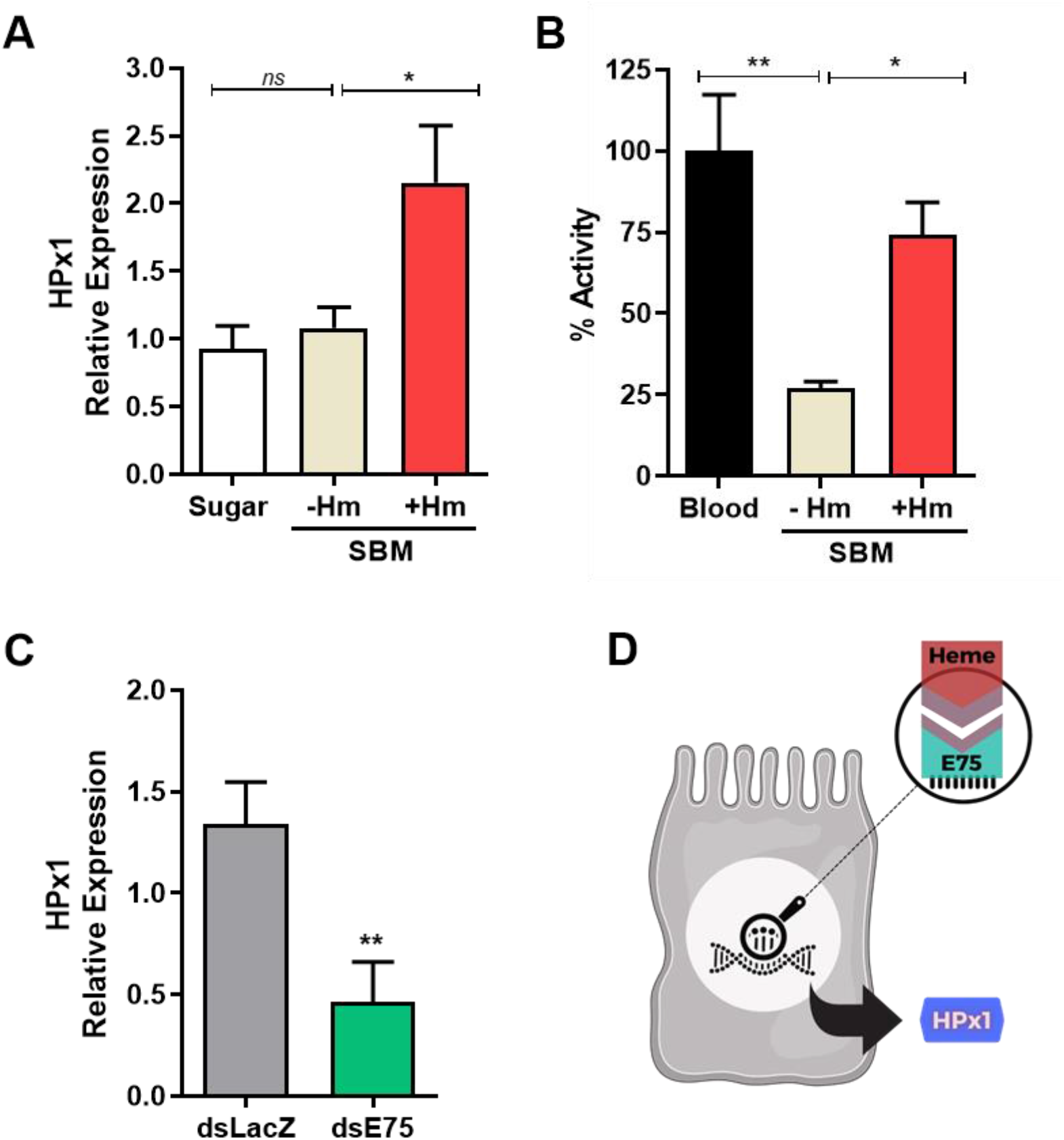
HPx1 expression is controlled by dietary heme and the E75 transcription factor. A) HPx1 expression in the midgut of sugar-fed or at 24 h after SBM feeding (without heme or supplemented with 50 μM of heme) (Sugar n=5; -Hm n=14; +Hm n=14). B) Catalytic activity of PMs from mosquitoes fed different diets at 24 h postfeeding (Blood n=3 pools of 10 PM each; -Hm n=4 pools of 10 PM each; +Hm n=4 pools of 10 PM each). C) HPx1 expression in midguts from control (dsLacZ) and dsE75-injected mosquitoes at 24 h ABM (dsLacZ n=3; dsE75 n=4). D) Schematic model of molecular signaling for HPx1 expression in the mosquito midgut. *p<0.05, **p<0.005, ns = not significant. Data are the mean +/-SEM. One-way ANOVA with Dunnett’s post-test for A and B and the T test for C.

### HPx1 contributes to PM assembly and regulation of the gut bacterial population

The PM is a semipermeable matrix that controls traffic of molecules between the intestinal lumen and the epithelia, and its correct assembly is essential to fulfilling its barrier function. Fig. 3A shows that control mosquitoes fed fluorescent dextran retained the polymer on the gut luminal side, a proxy of the barrier function of the PM. In contrast, HPx1-silenced mosquitoes presented strong fluorescence in the epithelial layer, suggesting a role for HPx1 in the proper assembly of the PM, as its permeability barrier function was compromised by HPx1 silencing. As this alteration in permeability might expose the epithelium to bacterial elicitors from the proliferative microbiota, we evaluated ROS production, known as an antimicrobial defense. Figs. 3B and 3B’ show that HPx1 silencing increased ROS levels in the midgut. The increase in ROS in HPx1-silenced mosquitoes was due to increased contact of the gut epithelia with the microbiota because oral administration of the antibiotics prevented the increase in the ROS levels, as observed in HPx1-silenced mosquitos (Fig. 3C). Exposure to damage signals and elevated ROS has been shown to activate intestinal stem cell mitosis (30). Indeed, HPx1 silencing increased phosphorylated H3-histone levels (Fig. 3D) in midgut epithelial cells, indicative of mitotic activity and suggestive of epithelial remodeling in response to oxidative imbalance. This highlights the key role of HPx1 in tissue homeostasis. The native midgut bacterial load was significantly reduced after HPx1 silencing (Fig. 3E). However, this was not due to canonical immune signaling pathways, as neither expression of two antimicrobial peptides, Attacin and Cecropin G, nor the bacterial sensor PGRP-LB by midgut cells was significantly different from that of dsLacZ controls (Fig. 3F), suggesting that ROS levels control microbiota proliferation.

**Figure 3.**
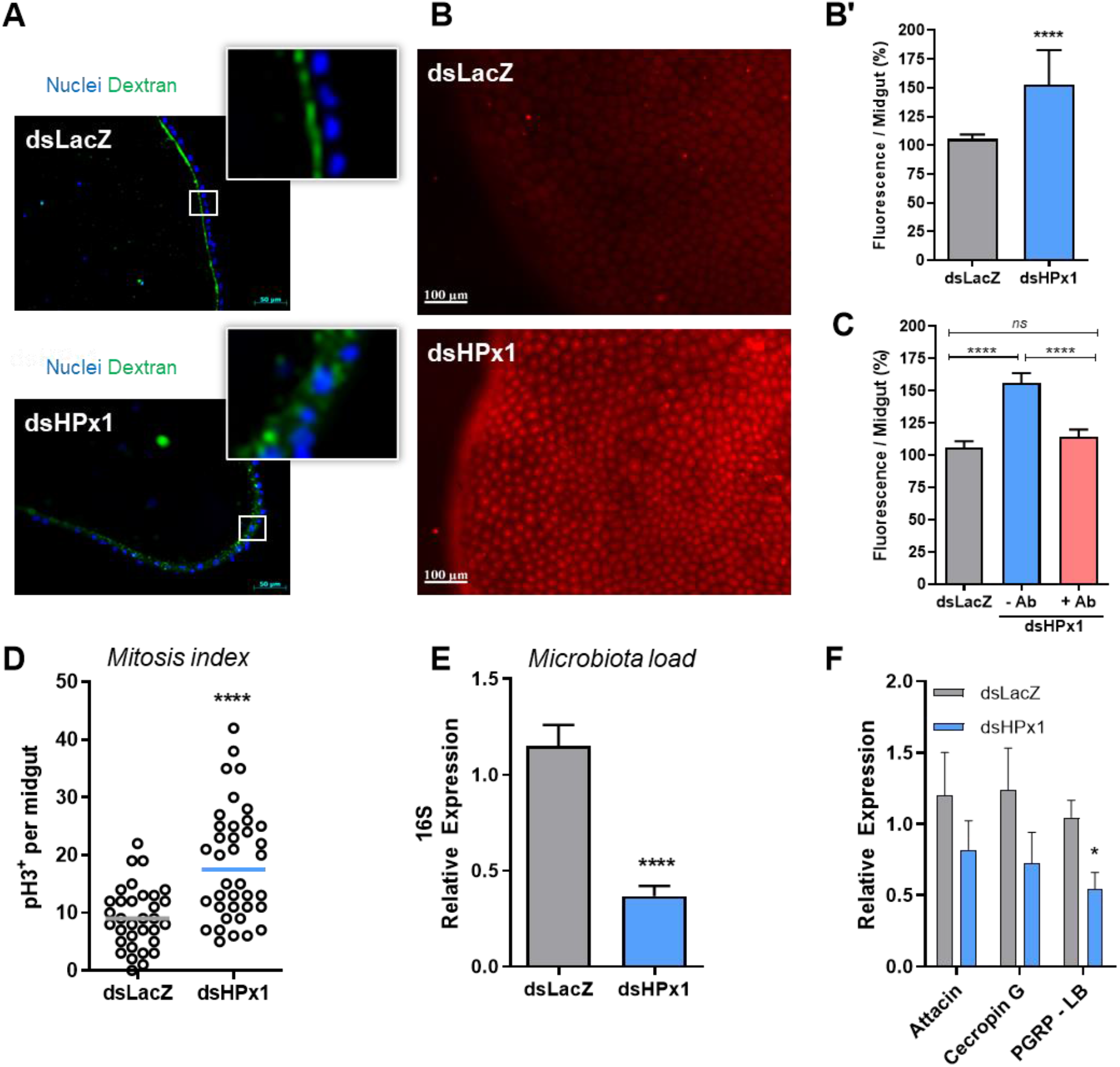
The role of HPx1 in PM assembly. A) Midgut transverse slices at 18 h ABM supplemented with dextran. Green: dextran – FITC. Blue: DAPI nuclear staining. Insets highlight dextran localization. B) Representative images of ROS levels measured by DHE oxidation in individual midguts at 18 h ABM. B’) Quantitative analysis of the fluorescence intensity of oxidized DHE (dsLacZ, n=35; dsHPx1, n=39). C) Quantitative analysis of the fluorescence intensity of oxidized DHE from individual midguts at 18 h ABM (dsLacZ, n=32; dsHPx1 - Ab, n=34; dsHPx1 + Ab, n=27). D) Mitosis index in the mosquito midgut at 18 h ABM (phospho-histone H3) (dsLacZ n=33; dsHPx1 n=38). E) The intestinal microbiota load analyzed through eubacterial ribosomal 16S gene expression by qPCR at 24 h ABM (dsLacZ n=6; dsHPx1 n=7). F) Immune-related gene expression upon HPx1 silencing at 24 h ABM by qPCR (dsLacZ n=8; dsHPx1 n=7). *p<0.05, ****p<0.0001, ns = not significant. Data are the mean +/-SEM. The T test for B’, D, E and F, and one-way ANOVA with Tukey’s posttest for C.

### HPx1 and DUOX coordinate intestinal immunity

NADPH-oxidases are a family of ROS-producing enzymes related to the immune system. Members of the dual oxidase (DUOX) group have been shown to play an essential role against bacterial challenge in the insect intestinal environment (Ha, Oh, Bae, et al., 2005; Kumar et al., 2010; Oliveira et al., 2011). Silencing HPx1 alone significantly increased ROS levels (Fig. 4A). In contrast, ROS levels were similar to those of dsLacZ controls in females in which HPx1 and DUOX were co-silenced (Fig. 4A and 4B), indicating that DUOX activity is the source of ROS when HPx1 is silenced. Unexpectedly, despite lowered ROS levels in double-silenced mosquitoes, bacterial levels remained reduced (Fig. 4C). This antibacterial response appears to be mediated by activation of canonical immune signaling pathways, as evidenced by increased expression of antimicrobial peptides and PGRP-LB in HPx1/DUOX co-silenced females (Fig. 4D-F).

**Figure 4.**
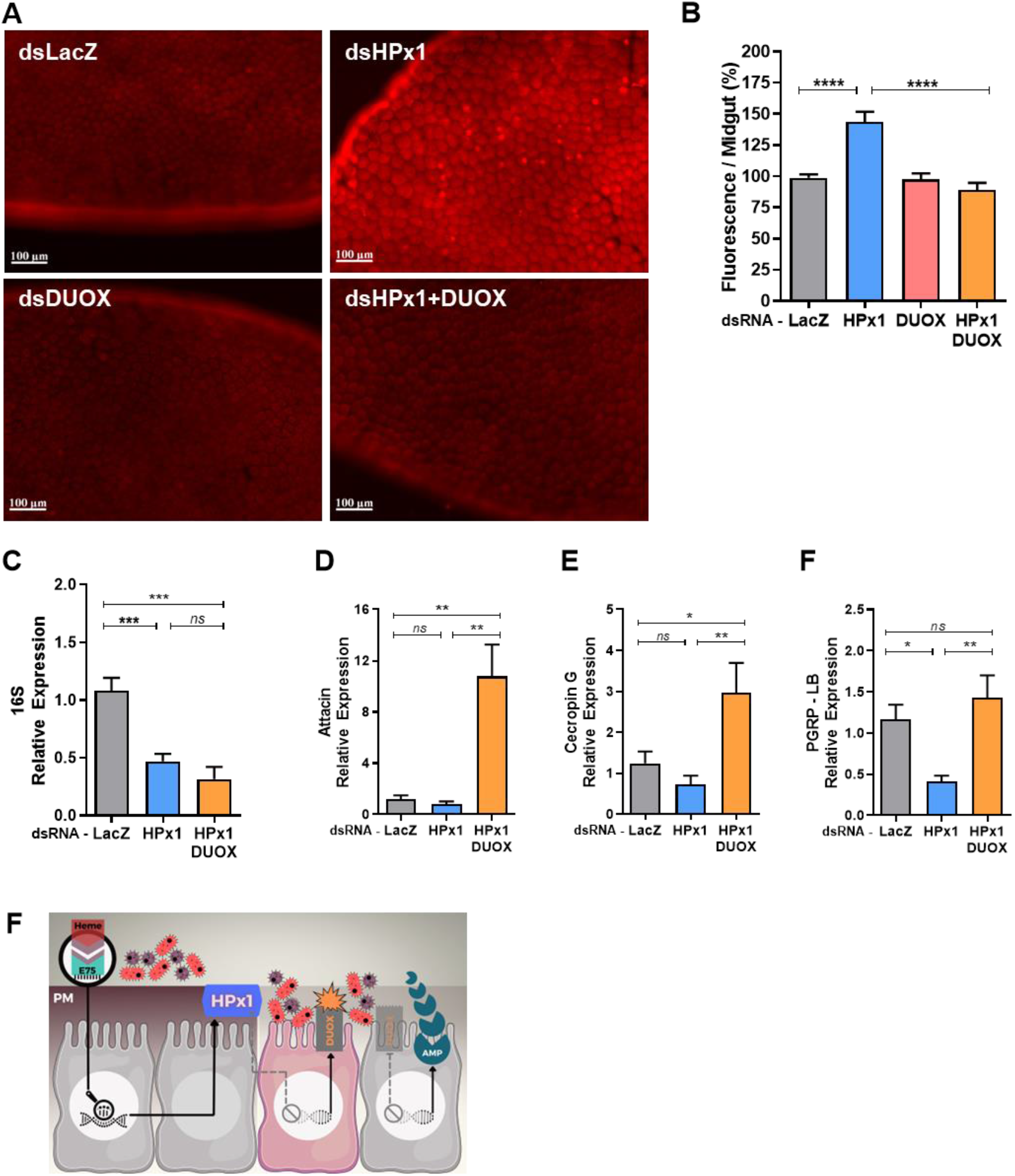
DUOX activation upon HPx1 silencing and hierarchical mode of activation of innate immunity in the mosquito midgut. A) Representative images of ROS levels measured by DHE oxidation in individual midguts at 18 h ABM. B) Quantitative analysis of the fluorescence intensity of oxidized DHE (dsLacZ, n=38; dsHPx1, n=43; dsDUOX, n=21; dsHPx1+DUOX, n=31). C) The intestinal microbiota load analyzed through eubacterial ribosomal 16S gene expression by qPCR at 24 h ABM (dsLacZ n=6; dsHPx1 n=6; dsHPx1+DUOX n=4). D-F) Immune-related gene expression upon HPx1 silencing at 24 h ABM by qPCR (at least n=5 for each condition). F) Schematic panel of intestinal immune activation showing that the PM integrity mediated by HPx1 activity isolates the gut microbiota, and once this integrity is lost, DUOX and antimicrobial peptides (AMPs) are activated. **p<0.005, ***p<0.001, ****p<0.0001, ns = not significant. Data are the mean +/-SEM. One-way ANOVA with Tukey’s post-test for A’, B, and C.

### Intestinal homeostasis impacts Zika and Dengue virus infection

The midgut epithelium is the first tissue that a virus must invade to establish a successful infection. By modulating diffusion of immune elicitors, HPx1 is crucial to maintaining gut homeostasis, allowing the proliferation of bacteria from the gut microbiota without triggering an immune response. Investigation of the effect of HPx1 silencing on midgut infection with Zika (ZKV) and Dengue virus (DENV) showed that silencing HPx1 expression dramatically reduced ZKV and DENV midgut infection at 4 and 7 days postfeeding, respectively (Fig. 5A and B). Titers of ZKV and DENV decreased approximately 100-fold in the gut of HPx1-silenced mosquitoes, along with a reduction in infection prevalence.

**Figure 5.**
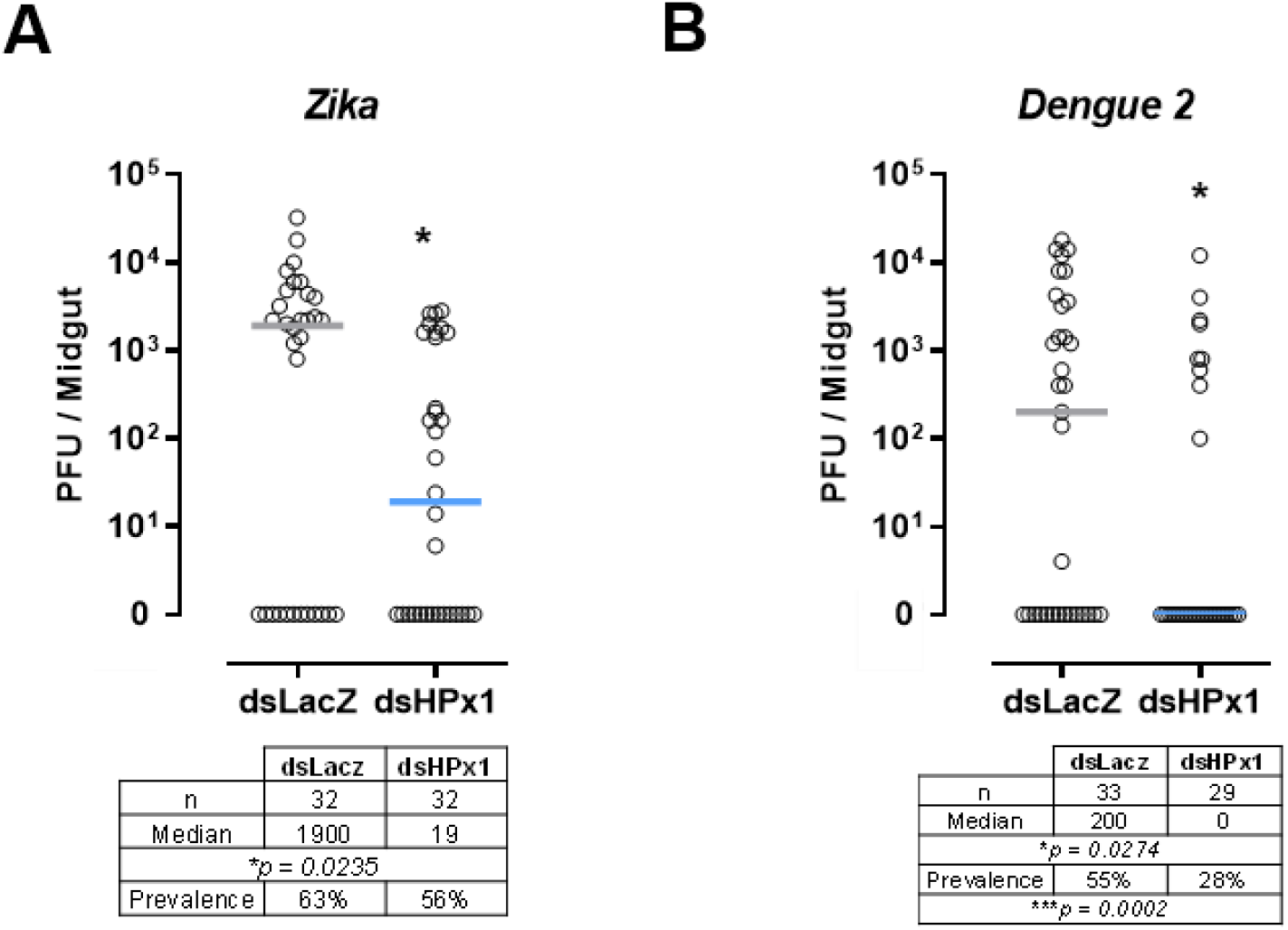
Viral infection in HPx1-silenced insects. A) Zika titers in midguts at 4 days postinfection. B) Dengue 2 titers in midguts at 7 days postinfection. Viral titers were assessed by the plaque assay. Each dot represents an individual mosquito gut, and bars indicate the median. * p< 0.05; Mann–Whitney test for A and B. The prevalence statistical analysis was performed by the Chi-square test followed by Fisher’s exact test.

## Discussion

The *Aedes aegypti* PM is an acellular layer that surrounds the blood bolus throughout the digestion process and limits direct contact of the epithelium with the midgut content and the intestinal microbiota, which undergoes massive proliferation upon blood feeding. In female mosquitoes, PM formation occurs in response to ingestion of a blood meal, following a time course that is finely coordinated with the pace of blood digestion. However, the signaling pathways that trigger PM secretion in adult mosquito females have not been elucidated nor has the impact of this structure on viral infection. Here, we characterize an intestinal secreted peroxidase (HPx1) that functions in PM assembly, contributing to its barrier function and promoting microbiota growth by preventing an antimicrobial response. This PM function has a permissive role for viral replication in the mosquito gut, thus constituting a novel determinant of vector competence. Importantly, dietary heme triggers HPx1 gene expression using the heme-dependent transcription factor E75, allowing synchronization of PM maturation with blood digestion by sensing the free heme released as hemoglobin is digested.

Similar to all blood-feeding organisms, mosquitoes face an oxidative challenge due to large amounts of heme – a pro-oxidant molecule – released by hemoglobin degradation. Therefore, preventing oxidative damage through ROS detoxification is a hallmark of their physiology (Sterkel et al., 2017). HPx1 mediates a novel mechanism to promote redox balance in the *Aedes* midgut through its hydrogen peroxide scavenging activity and by modulating PM barrier function. We also show that HPx1 allows proliferation of the gut microbiota without activating a DUOX-mediated oxidative burst by limiting exposure of gut epithelial cells to microbial immune elicitors. We have previously shown that *Aedes aegypti* catalase (AAEL013407-RB), the main intracellular hydrogen peroxide scavenger, is induced in the blood-fed midgut of females (Oliveira et al., 2017). Here, we demonstrate that HPx1 contributes to the overall peroxide scavenging capacity of the gut in a way that is independent of epithelial intracellular catalase but at comparable activity levels (Fig 1). Although peroxidases are less efficient in decomposing hydrogen peroxide than catalases, it is known that some heme peroxidases also have high catalase activity (Vidossich et al., 2012; Vlasits et al., 2010).

*A. gambiae* IMPer/HPx15 belongs to anopheline-specific expansion of the heme peroxidase family (Kajla et al., 2016) nested in the same branch of the peroxidase family of *A. aegypti* HPx1 and immune-regulated catalase (IRC; CG8913) of *D. melanogaster* (Ha, Oh, Ryu, et al., 2005; Konstandi et al., 2005; Waterhouse et al., 2007). However, *Drosophila* IRC has a conserved heme peroxidase domain structure (Pfam PF03098) but lacks a catalase domain (Pfam PF00199), despite having a high hydrogen peroxide dismutation activity (Ha, Oh, Ryu, et al., 2005), which is a feature of catalases but uncommon among peroxidases. Interestingly, IMPer/HPx15 of *Anopheles* is also expressed in the female reproductive tract, induced by the ecdysone transferred along with the sperm during insemination, due to ecdysone-responsive elements present in the promoter region (Shaw et al., 2014). Ecdysone-responsive element sequences close to the *Aedes* HPx1 gene have been identified *in silico* (Zhao et al., 2001). Therefore, *Aedes* HPx1, IRC, and AgIMPer are secreted enzymes that modulate interactions of the midgut with commensal and pathogenic bacteria. Nevertheless, their role in the biology of the reproductive organs has not been well established. Although our data reveal that HPx1, IRC, and IMPer share sequence and functional homology, it is not possible at present to speculate which roles are ancestral and which were acquired secondarily during the evolution of dipterans.

*A. gambiae* IMPer has been proposed to crosslink external matrix proteins by forming di-tyrosine bridges, reducing the accessibility of microbial elicitors to the intestinal cells (Kumar et al., 2010). In *Drosophila*, however, a transglutaminase enzyme crosslinks PM proteins, also protecting the midgut epithelia from damage (Kuraishi et al., 2011; Shibata et al., 2015). In this study, we show that HPx1 associates with the PM and modulates gut permeability in *A. aegypti* (Fig. 4). When the PM structure was compromised by HPx1 silencing, we observed immediate responses from the epithelium that increased ROS levels, which was attributed mainly to DUOX activation by microbial elicitors. The role of PM in preventing elevated ROS production in the gut epithelium was also observed when the PM was compromised by the chitin inhibitor diflubenzuron administered in a blood meal for *A. aegypti* females (Taracena et al., 2018). A simple hypothesis to explain how elevated ROS might lower virus infection is that they directly attack the virus. However, arbovirus infection of midgut cells is thought to occur early during digestion, several hours before proliferation of microbiota occurs and before the PM is secreted (Franz et al., 2015). Therefore, it is unlikely that extracellular ROS produced in response to bacterial elicitors in HPx1-silenced insects would directly attack virus particles that will be localized intracellularly when these molecules start to increase in the lumen. In general, cellular antiviral mechanisms are most plausibly responsible for hampering viral infection upon HPx1 silencing. Among these possible mechanisms, the elevated extracellular ROS levels derived from DUOX activation are indeed sensed by the gut as a danger signal, evoking a tissue-repairing response the hallmark of which is stem cell proliferation (Fig. 4), a homeostatic response coupled to cell death in response to insult and damage. Interestingly, Taracena et al. proposed that different degrees of resistance to infection among mosquito strains are related to different capacities to promote a rapid increase in stem cell proliferation; hence, faster cell death, followed by cell renewal from stem cell activation, is a process that reduces viral infection (Taracena et al., 2018). One could hypothesize that this mechanism is responsible for the decrease in ZKV and DENV infection promoted by HPx1 silencing.

In mammals, the mucus layer allows commensal bacteria from the gut microbiota to thrive without eliciting microbicidal immune responses by the intestinal mucosa. In other words, the mammalian mucus layer acts as a physical barrier that leads to “immunological ignorance” by preventing a state of constant immune activation and chronic inflammation of the intestine in response to immune elicitors from the normal microbiota (Chassaing et al., 2015; Hooper, 2009; Macpherson et al., 2005). In mosquitos, the PM is secreted when the microbiota peaks in number to approximately 100-1000 times the population found before a blood meal (Oliveira et al., 2011), representing a potential massive immune challenge to this tissue. Hixson et al. suggested that immune tolerance to the indigenous microbiota might be mediated by high expression of caudal and PGRP genes, leading to low expression of antimicrobial peptides in epithelial cells from the posterior gut (Hixson et al., n.d.). Here, we highlight the role of PM-associated HPx1 in limiting exposure of the epithelium to immune elicitors from the expanded microbiota observed postfeeding. Before blood feeding, the PM is absent, and bacteria interact with the epithelium, leading to ROS generation and damage-induced repair (Oliveira et al., 2011; Taracena et al., 2018). When the first line of immediate response to immune elicitors (redox mediated) is further prevented by simultaneous DUOX silencing, a reaction is activated (antibacterial peptides expression) to limit microbial growth. These data suggest hierarchal immune activation in the *A. aegypti* midgut fundamentally orchestrated by the PM integrity. This is similar to what was reported for *A. gambiae*, whereby bacterial elicitors lead to DUOX-mediated activation of IMPer, which promotes cross-linking of extracellular proteins (Kumar et al., 2010). However, in this report, an epithelial cell mucous layer (and not the PM) was indicated as the site of action of the peroxidase. Regarding the mode of activation of this pathway, the findings shown herein are also fairly similar to those for *Drosophila*, with ROS produced by DUOX being primarily triggered by bacterial pathogens, and the IMD pathway induces antimicrobial peptide production upon activation failure (Buchon et al., 2009; Ha et al., 2009; Hixson et al., n.d.; Ryu et al., 2006).

Traditionally, immunology has focused on how hosts eliminate pathogens while fighting infections, but in the last decade, there has been a growing interest in how hosts endure infection by utilizing disease tolerance, including diminishing both the direct damage caused by the pathogen and the self-inflicted damage due to the host immune reaction directed at the elimination of the pathogen. In this study, we showed that PM-associated HPx1 is pivotal to maintaining gut immune homeostasis, acting as a tolerance mechanism that prevents responses to the microbiota. Additionally, the fact that HPx1/PM disruption causes a drastic reduction in the viral load within the gut epithelia suggests that this mechanism is essential to the tolerogenic status of the gut to viral replication and directly contributes to vector competence.

Together, our results indicate that the *A. aegypti* PM supports midgut homeostasis during blood digestion. Heme derived from blood hemoglobin digestion regulates expression of HPx1, an enzyme that has a central role in the assembly of a fully functional PM, through the heme-sensitive E75 transcription factor. Thus, HPx1 maintains immunological ignorance of the midgut epithelia toward the microbiota, allowing a state of microbiota and viral tolerance and preventing tissue damage.

## Acknowledgments

We thank all members of the Laboratory of Biochemistry of Hematophagous Arthropods, especially Jaciara Miranda Freire, for rearing the insects and Patricia Ingridis S. Cavalcante, João Marques, Charlion Cosme and S.R. Cassia for providing technical assistance. This work was supported by grants from the Conselho Nacional de Desenvolvimento Científico e Tecnológico (CNPq), Coordenação de Aperfeiçoamento de Pessoal de Nível Superior (CAPES), Financiadora de Estudos e Projetos (FINEP) and Fundação de Amparo a Pesquisa do Estado do Rio de Janeiro (FAPERJ).

## Supplementary figures

**Supplementary Figure 1.**
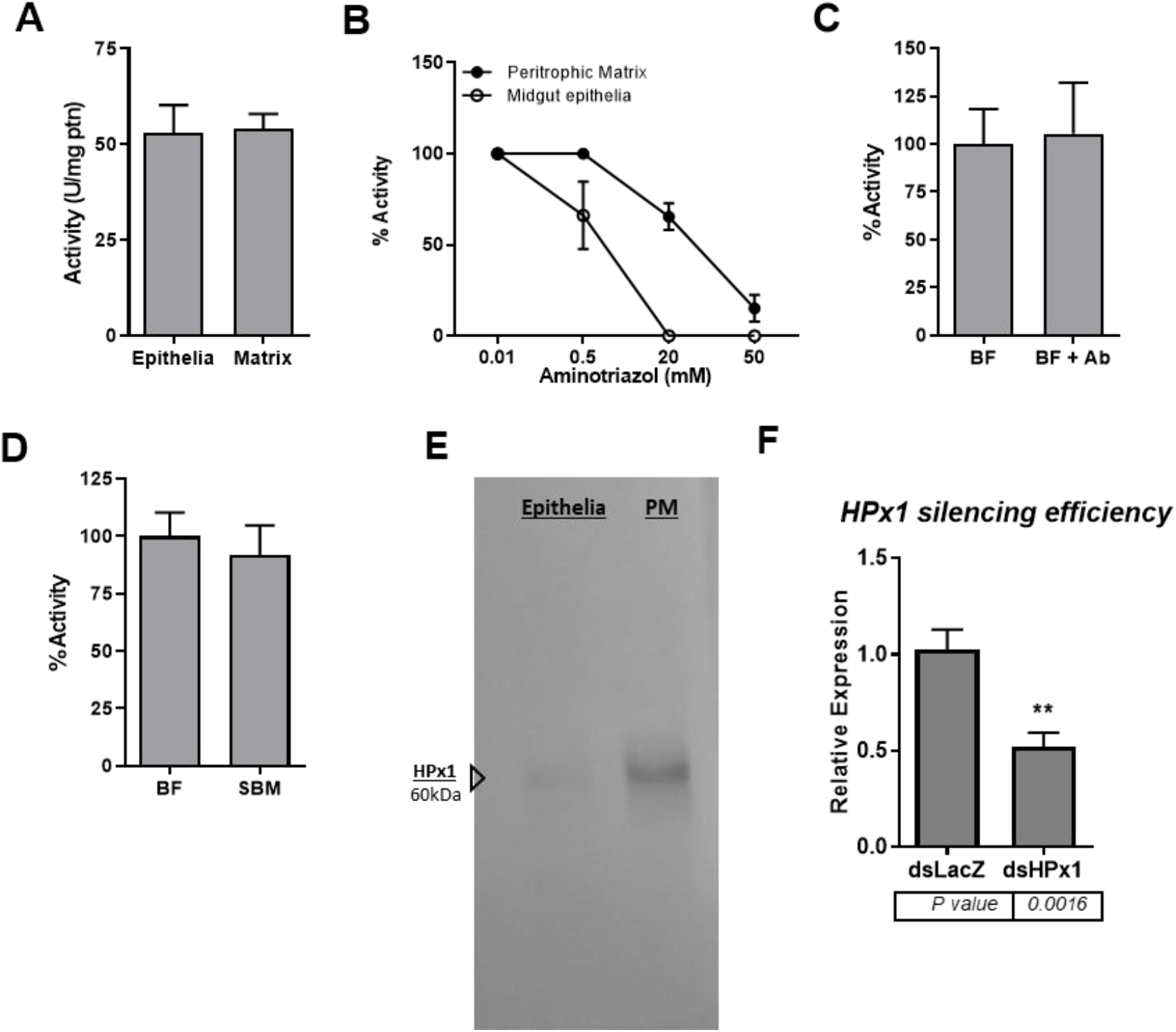
SUP1: A) Catalase-specific activity comparison of the intestinal epithelia and PM at 24 h ABM. B) *In vitro* sensitivity of midgut epithelia and PM samples to aminotriazole, a catalase/peroxidase inhibitor. C) Mosquitos were treated with (or without) an antibiotic cocktail in a sugar meal, and PM catalase activity was assayed at 24 h ABM (BF n=10; +AB n=11). (D) Catalase activity comparison of the PM at 24 h ABM for mosquitoes fed blood or SBM, a chemically defined artificial diet (BF n=21; +AB n=17). E) HPx1 western blot of midgut epithelia and PM protein extracts, referring to Fig. 1E.

**Supplementary Figure 2.**
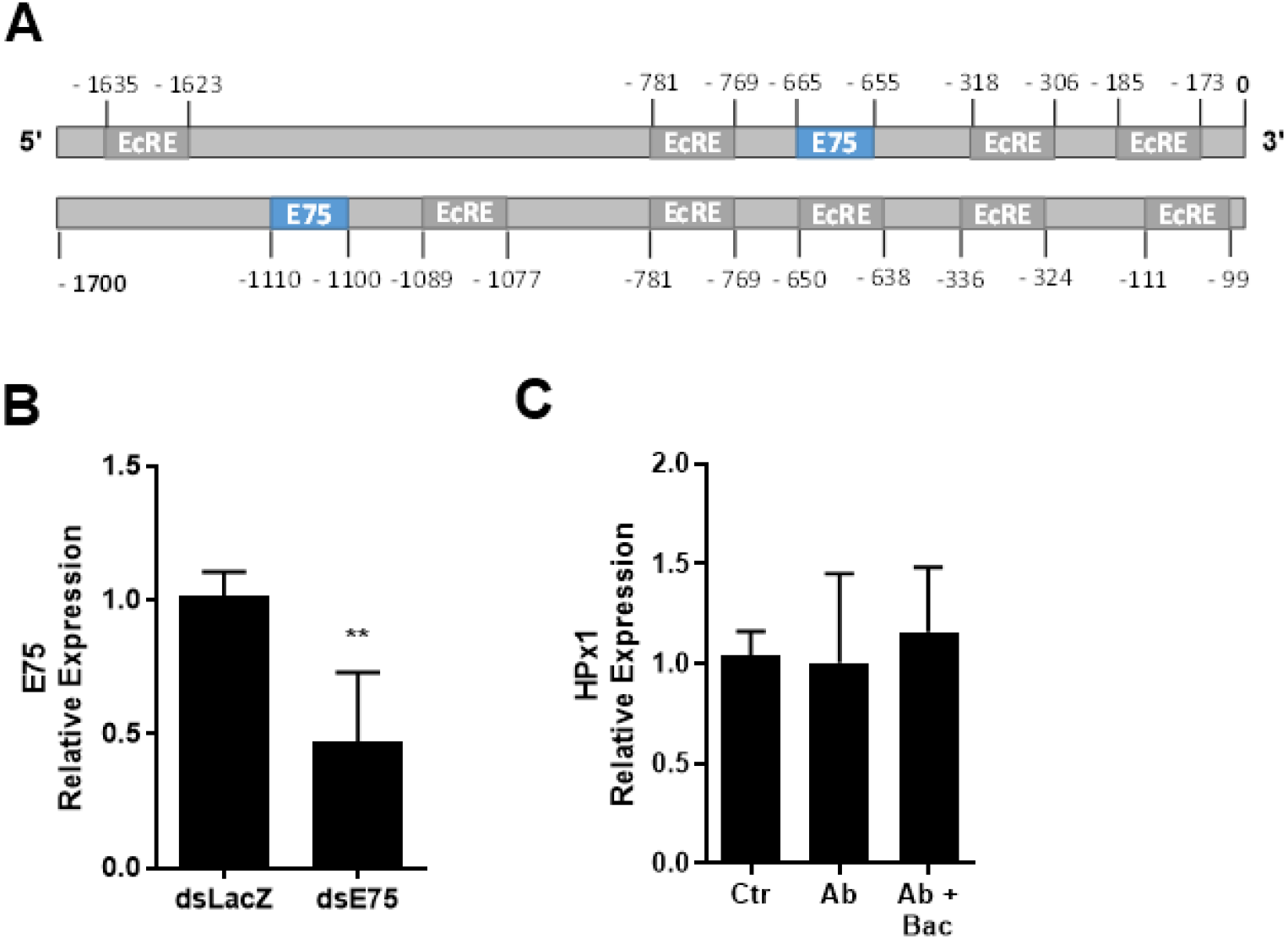
SUP2: A) Schematic illustration of the E75 and ecdysone receptor-binding motifs in the promoter region of HPx1. Numbers refer to nucleotide positions relative to the transcription start site. B) E75 silencing efficiency in midguts at 24 h ABM. C) HPx1 expression in midguts at 24 h ABM. Control mosquitoes were fed a regular sucrose solution before blood feeding. Ab mosquitoes were pretreated with antibiotics before blood feeding. Ab + Bac mosquitoes were pretreated with antibiotics and fed blood containing *Enterobacter cloacae at* 1 OD/ml (Ctr n=6; Ab n=6; Ab+Bac n=5). Data are the mean +/-SEM.

**Supplementary Table 1.**
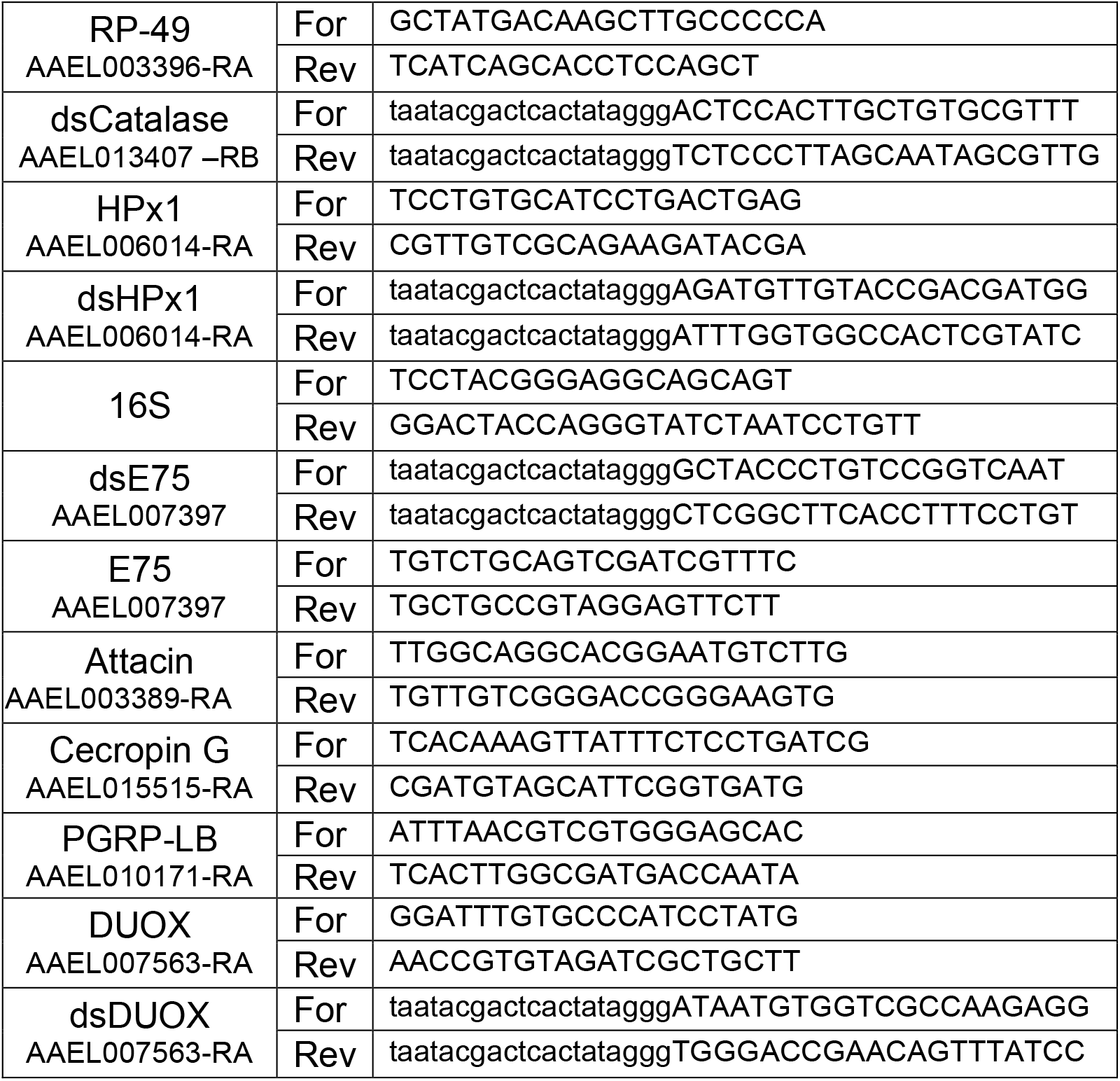
Primer list.

## References

Aebi, H. (1984). Catalase in vitro. In Methods in Enzymology (Vol. 105, Issue 1947, pp. 121–126). https://doi.org/10.1016/S0076-6879(84)05016-3

Bailey, T. L., Boden, M., Buske, F. A., Frith, M., Grant, C. E., Clementi, L., Ren, J., Li, W. W., & Noble, W. S. (2009). MEME SUITE: tools for motif discovery and searching. Nucleic Acids Research, 37(Web Server), W202–W208. https://doi.org/10.1093/nar/gkp335

Black IV, W. C., Bennett, K. E., Gorrochótegui-Escalante, N., Barillas-Mury, C. v., Fernández-Salas, I., Muñoz, M. D. L., Farfán-Alé, J. A., Olson, K. E., & Beaty, B. J. (2002). Flavivirus susceptibility in Aedes aegypti. Archives of Medical Research, 33(4), 379–388. https://doi.org/10.1016/S0188-4409(02)00373-9

Bottino-Rojas, V., Talyuli, O. A. C., Carrara, L., Martins, A. J., James, A. A., Oliveira, P. L., & Paiva-Silva, G. O. (2018). The redox-sensing gene Nrf2 affects intestinal homeostasis, insecticide resistance, and Zika virus susceptibility in the mosquito Aedes aegypti. Journal of Biological Chemistry, 293(23), 9053–9063. https://doi.org/10.1074/jbc.RA117.001589

Bottino-Rojas, V., Talyuli, O. A. C., Jupatanakul, N., Sim, S., Dimopoulos, G., Venancio, T. M., Bahia, A. C., Sorgine, M. H., Oliveira, P. L., & Paiva-Silva, G. O. (2015). Heme signaling impacts global gene expression, immunity and dengue virus infectivity in Aedes aegypti. PLoS ONE, 10(8). https://doi.org/10.1371/journal.pone.0135985

Buchon, N., Broderick, N. a, Poidevin, M., Pradervand, S., & Lemaitre, B. (2009). Drosophila intestinal response to bacterial infection: activation of host defense and stem cell proliferation. Cell Host & Microbe, 5(2), 200–211. https://doi.org/10.1016/j.chom.2009.01.003

Chassaing, B., Koren, O., Goodrich, J. K., Poole, A. C., Srinivasan, S., Ley, R. E., & Gewirtz, A. T. (2015). Dietary emulsifiers impact the mouse gut microbiota promoting colitis and metabolic syndrome. Nature, 519(7541), 92–96. https://doi.org/10.1038/nature14232

Coutinho-Abreu, I. v., Sharma, N. K., Robles-Murguia, M., & Ramalho-Ortigao, M. (2010). Targeting the Midgut Secreted PpChit1 Reduces Leishmania major Development in Its Natural Vector, the Sand Fly Phlebotomus papatasi. PLoS Neglected Tropical Diseases, 4(11), e901. https://doi.org/10.1371/journal.pntd.0000901

Cruz, J., Mane-Padros, D., Zou, Z., & Raikhel, A. S. (2012). Distinct roles of isoforms of the heme-liganded nuclear receptor E75, an insect ortholog of the vertebrate Rev-erb, in mosquito reproduction. Molecular and Cellular Endocrinology, 349(2), 262–271. https://doi.org/10.1016/j.mce.2011.11.006

Devenport, M., Alvarenga, P. H., Shao, L., Fujioka, H., Bianconi, M. L., Oliveira, P. L., & Jacobs-Lorena, M. (2006). Identification of the Aedes aegypti peritrophic matrix protein AeIMUCI as a heme-binding protein. Biochemistry, 45(31), 9540–9549. https://doi.org/10.1021/bi0605991

Franz, A., Kantor, A., Passarelli, A., & Clem, R. (2015). Tissue Barriers to Arbovirus Infection in Mosquitoes. Viruses, 7(7), 3741–3767. https://doi.org/10.3390/v7072795

Garver, L. S., de Almeida Oliveira, G., & Barillas-Mury, C. (2013). The JNK Pathway Is a Key Mediator of Anopheles gambiae Antiplasmodial Immunity. PLoS Pathogens, 9(9), e1003622. https://doi.org/10.1371/journal.ppat.1003622

Gentile, C., Lima, J., & Peixoto, A. (2005). Isolation of a fragment homologous to the rp49 constitutive gene of Drosophila in the Neotropical malaria vector Anopheles aquasalis (Diptera: Culicidae). Memórias Do Instituto Oswaldo Cruz, 100(October), 545–547. http://www.scielo.br/scielo.php?pid=S0074-02762005000600008&script=sci_arttext

Grant, C. E., Bailey, T. L., & Noble, W. S. (2011). FIMO: scanning for occurrences of a given motif. Bioinformatics, 27(7), 1017–1018. https://doi.org/10.1093/bioinformatics/btr064

Ha, E.-M., Lee, K.-A., Seo, Y. Y., Kim, S.-H., Lim, J.-H., Oh, B.-H., Kim, J., & Lee, W.-J. (2009). Coordination of multiple dual oxidase-regulatory pathways in responses to commensal and infectious microbes in drosophila gut. Nature Immunology, 10(9), 949–957. https://doi.org/10.1038/ni.1765

Ha, E.-M., Oh, C.-T., Bae, Y. S., & Lee, W.-J. (2005). A direct role for dual oxidase in Drosophila gut immunity. Science (New York, N.Y.), 310(5749), 847–850. https://doi.org/10.1126/science.1117311

Ha, E.-M., Oh, C.-T., Ryu, J.-H., Bae, Y.-S., Kang, S.-W., Jang, I.-H., Brey, P. T., & Lee, W.-J. (2005). An Antioxidant System Required for Host Protection against Gut Infection in Drosophila. Developmental Cell, 8(1), 125–132. https://doi.org/10.1016/j.devcel.2004.11.007

Hixson, B., Bing, X.-L., Yang, X., Bonfini, A., & Nagy, P. (n.d.). A transcriptomic atlas of Aedes aegypti reveals detailed functional organization of major body parts and gut regional specializations in sugar-fed and blood-fed adult females. https://doi.org/10.1101/2021.12.19.473372

Hooper, L. v. (2009). Do symbiotic bacteria subvert host immunity? Nature Reviews. Microbiology, 7(5), 367–374. https://doi.org/10.1038/nrmicro2114

Jaramillo-Gutierrez, G., Molina-Cruz, A., Kumar, S., & Barillas-Mury, C. (2010). The Anopheles gambiae Oxidation Resistance 1 (OXR1) Gene Regulates Expression of Enzymes That Detoxify Reactive Oxygen Species. PLoS ONE, 5(6), e11168. https://doi.org/10.1371/journal.pone.0011168

Jupatanakul, N., Sim, S., Angleró-Rodríguez, Y. I., Souza-Neto, J., Das, S., Poti, K. E., Rossi, S. L., Bergren, N., Vasilakis, N., & Dimopoulos, G. (2017). Engineered Aedes aegypti JAK/STAT Pathway-Mediated Immunity to Dengue Virus. PLOS Neglected Tropical Diseases, 11(1), e0005187. https://doi.org/10.1371/journal.pntd.0005187

Kajla, M., Biol, J. P. E., Kajla, M., Gupta, K., Kakani, P., Dhawan, R., Choudhury, T. P., Gupta, L., & Gakhar, S. K. (2016). Identification of an Anopheles Lineage-Specific Unique Heme Peroxidase HPX15: A Plausible Candidate for Arresting Malaria Parasite Development. 3(4). https://doi.org/10.4172/2329-9002.1000160

Kokoza, V. A., Martin, D., Mienaltowski, M. J., Ahmed, A., Morton, C. M., & Raikhel, A. S. (2001). Transcriptional regulation of the mosquito vitellogenin gene via a blood meal-triggered cascade. Gene, 274(1–2), 47–65. https://doi.org/10.1016/S0378-1119(01)00602-3

Konstandi, O. A., Papassideri, I. S., Stravopodis, D. J., Kenoutis, C. A., Hasan, Z., Katsorchis, T., Wever, R., & Margaritis, L. H. (2005). The enzymatic component of Drosophila melanogaster chorion is the Pxd peroxidase. Insect Biochemistry and Molecular Biology, 35(9), 1043–1057. https://doi.org/10.1016/j.ibmb.2005.04.005

Kumar, S., & Barillas-Mury, C. (2005). Ookinete-induced midgut peroxidases detonate the time bomb in anopheline mosquitoes. Insect Biochemistry and Molecular Biology, 35(7), 721–727. https://doi.org/10.1016/j.ibmb.2005.02.014

Kumar, S., Molina-Cruz, A., Gupta, L., Rodrigues, J., & Barillas-Mury, C. (2010). A Peroxidase/Dual Oxidase System Modulates Midgut Epithelial Immunity in Anopheles gambiae. Science, 327(5973), 1644–1648. https://doi.org/10.1126/science.1184008

Kuraishi, T., Binggeli, O., Opota, O., Buchon, N., & Lemaitre, B. (2011). Genetic evidence for a protective role of the peritrophic matrix against intestinal bacterial infection in Drosophila melanogaster. Proceedings of the National Academy of Sciences, 108(38), 15966–15971. https://doi.org/10.1073/pnas.1105994108

Kuraishi, T., Hori, A., & Kurata, S. (2013). Host-microbe interactions in the gut of Drosophila melanogaster. Frontiers in Physiology, 4(December), 1–8. https://doi.org/10.3389/fphys.2013.00375

Lehane, M. J. (1997). Peritrophic Matrix Structure and Function. Annual Review of Entomology, 42(1), 525–550. https://doi.org/10.1146/annurev.ento.42.1.525

Liu, J., Liu, Y., Nie, K., Du, S., Qiu, J., Pang, X., Wang, P., & Cheng, G. (2016). Flavivirus NS1 protein in infected host sera enhances viral acquisition by mosquitoes. Nature Microbiology, 1(9), 16087. https://doi.org/10.1038/nmicrobiol.2016.87

Lowry, O. H., Rosebrough, N. J., Farr, A. L., & Randall, R. J. (1951). Protein measurement with the Folin phenol reagent. The Journal of Biological Chemistry, 193(1), 265–275. http://linkinghub.elsevier.com/retrieve/pii/S0003269784711122

Macpherson, A. J., Geuking, M. B., & McCoy, K. D. (2005). Immune responses that adapt the intestinal mucosa to commensal intestinal bacteria. Immunology, 115(2), 153–162. https://doi.org/10.1111/j.1365-2567.2005.02159.x

Oliveira, J. H. M., Gonçalves, R. L. S., Lara, F. A., Dias, F. A., Gandara, A. C. P., Menna-Barreto, R. F. S., Edwards, M. C., Laurindo, F. R. M., Silva-Neto, M. a C., Sorgine, M. H. F., & Oliveira, P. L. (2011). Blood meal-derived heme decreases ROS levels in the midgut of Aedes aegypti and allows proliferation of intestinal microbiota. PLoS Pathogens, 7(3), e1001320. https://doi.org/10.1371/journal.ppat.1001320

Oliveira, J. H. M., Talyuli, O. A. C., Goncalves, R. L. S., Paiva-Silva, G. O., Sorgine, M. H. F., Alvarenga, P. H., & Oliveira, P. L. (2017). Catalase protects Aedes aegypti from oxidative stress and increases midgut infection prevalence of Dengue but not Zika. PLOS Neglected Tropical Diseases, 11(4), e0005525. https://doi.org/10.1371/journal.pntd.0005525

Pascoa, V., Oliveira, P. L., Dansa-Petretski, M., Silva, J. R., Alvarenga, P. H., Jacobs-Lorena, M., & Lemos, F. J. A. (2002). Aedes aegypti peritrophic matrix and its interaction with heme during blood digestion. Insect Biochemistry and Molecular Biology, 32(5), 517–523. https://doi.org/10.1016/S0965-1748(01)00130-8

Petersen, T. N., Brunak, S., von Heijne, G., & Nielsen, H. (2011). SignalP 4.0: discriminating signal peptides from transmembrane regions. Nature Methods, 8(10), 785–786. https://doi.org/10.1038/nmeth.1701

Pfaffl, M. W. (2001). A new mathematical model for relative quantification in real-time RT–PCR. Nucleic Acids Research, 29(9), e45. http://www.pubmedcentral.nih.gov/articlerender.fcgi?artid=55695&tool=pmcentrez&rendertype=abstract

Ramalho-Ortigao, M. (2010). Sand Fly-Leishmania Interactions: Long Relationships are Not Necessarily Easy. The Open Parasitology Journal, 4(1), 195–204. https://doi.org/10.2174/1874421401004010195

Retamal, C. A., Thiebaut, P., & Alves, E. W. (1999). Protein Purification from Polyacrylamide Gels by Sonication Extraction. Analytical Biochemistry, 268(1), 15–20. https://doi.org/10.1006/abio.1998.2977

Rose, C., Casas-Sánchez, A., Dyer, N. A., Solórzano, C., Beckett, A. J., Middlehurst, B., Marcello, M., Haines, L. R., Lisack, J., Engstler, M., Lehane, M. J., Prior, I. A., & Acosta-Serrano, Á. (2020). Trypanosoma brucei colonizes the tsetse gut via an immature peritrophic matrix in the proventriculus. Nature Microbiology, 2. https://doi.org/10.1038/s41564-020-0707-z

Ryu, J.-H., Ha, E.-M., Oh, C.-T., Seol, J.-H., Brey, P. T., Jin, I., Lee, D. G., Kim, J., Lee, D., & Lee, W.-J. (2006). An essential complementary role of NF-κB pathway to microbicidal oxidants in Drosophila gut immunity. The EMBO Journal, 25(15), 3693–3701. https://doi.org/10.1038/sj.emboj.7601233

Shahabuddin, M., Kaidoh, T., Aikawa, M., & Kaslow, D. C. (1995). Plasmodium gallinaceum: Mosquito peritrophic matrix and the parasite-vector compatibility. Experimental Parasitology, 81(3), 386–393. https://doi.org/10.1006/expr.1995.1129

Shao, L., Devenport, M., & Jacobs-lorena, M. (2001). The Peritrophic Matrix of Hematophagous Insects. Archives of Insect Biochemistry and Physiology, 47(November 2000), 119–125. https://doi.org/10.1002/arch.1042

Shaw, W. R., Teodori, E., Mitchell, S. N., Baldini, F., Gabrieli, P., Rogers, D. W., & Catteruccia, F. (2014). Mating activates the heme peroxidase HPX15 in the sperm storage organ to ensure fertility in Anopheles gambiae. Proceedings of the National Academy of Sciences, 111(16), 5854–5859. https://doi.org/10.1073/pnas.1401715111

Shibata, T., Maki, K., Hadano, J., Fujikawa, T., Kitazaki, K., Koshiba, T., & Kawabata, S. I. (2015). Crosslinking of a Peritrophic Matrix Protein Protects Gut Epithelia from Bacterial Exotoxins. PLoS Pathogens, 11(10), 1–15. https://doi.org/10.1371/journal.ppat.1005244

Sterkel, M., Oliveira, J. H. M., Bottino-Rojas, V., Paiva-Silva, G. O., & Oliveira, P. L. (2017). The Dose Makes the Poison: Nutritional Overload Determines the Life Traits of Blood-Feeding Arthropods. Trends in Parasitology, 33(8), 633–644. https://doi.org/10.1016/j.pt.2017.04.008

Talyuli, O. A. C., Bottino-Rojas, V., Polycarpo, C. R., Oliveira, P. L., & Paiva-Silva, G. O. (2021). Non-immune Traits Triggered by Blood Intake Impact Vectorial Competence. In Frontiers in Physiology (Vol. 12). Frontiers Media S.A. https://doi.org/10.3389/fphys.2021.638033

Talyuli, O. A. C., Bottino-Rojas, V., Taracena, M. L., Soares, A. L. M., Oliveira, J. H. M., & Oliveira, P. L. (2015). The use of a chemically defined artificial diet as a tool to study Aedes aegypti physiology. Journal of Insect Physiology, 83, 1–7. https://doi.org/10.1016/j.jinsphys.2015.11.007

Taracena, M. L., Bottino-Rojas, V., Talyuli, O. A. C., Walter-Nuno, A. B., Oliveira, J. H. M., Angleró-Rodriguez, Y. I., Wells, M. B., Dimopoulos, G., Oliveira, P. L., & Paiva-Silva, G. O. (2018). Regulation of midgut cell proliferation impacts Aedes aegypti susceptibility to dengue virus. PLOS Neglected Tropical Diseases, 12(5), e0006498. https://doi.org/10.1371/journal.pntd.0006498

Terra, W. R., Dias, R. O., Oliveira, P. L., Ferreira, C., & Venancio, T. M. (2018). Transcriptomic analyses uncover emerging roles of mucins, lysosome/secretory addressing and detoxification pathways in insect midguts. Current Opinion in Insect Science, 29, 34–40. https://doi.org/10.1016/j.cois.2018.05.015

Vidossich, P., Alfonso-Prieto, M., & Rovira, C. (2012). Catalases versus peroxidases: DFT investigation of H2O2 oxidation in models systems and implications for heme protein engineering. Journal of Inorganic Biochemistry, 117, 292–297. https://doi.org/10.1016/j.jinorgbio.2012.07.002

Vlasits, J., Jakopitsch, C., Bernroitner, M., Zamocky, M., Furtmüller, P. G., & Obinger, C. (2010). Mechanisms of catalase activity of heme peroxidases. Archives of Biochemistry and Biophysics, 500(1), 74–81. https://doi.org/10.1016/j.abb.2010.04.018

Waterhouse, R. M., Kriventseva, E. v., Meister, S., Xi, Z., Alvarez, K. S., Bartholomay, L. C., Barillas-Mury, C., Bian, G., Blandin, S., Christensen, B. M., Dong, Y., Jiang, H., Kanost, M. R., Koutsos, A. C., Levashina, E. A., Li, J., Ligoxygakis, P., MacCallum, R. M., Mayhew, G. F., … Christophides, G. K. (2007). Evolutionary Dynamics of Immune-Related Genes and Pathways in Disease-Vector Mosquitoes. Science, 316(5832), 1738–1743. https://doi.org/10.1126/science.1139862

Weiss, B. L., Savage, A. F., Griffith, B. C., Wu, Y., & Aksoy, S. (2014). The Peritrophic Matrix Mediates Differential Infection Outcomes in the Tsetse Fly Gut following Challenge with Commensal, Pathogenic, and Parasitic Microbes. The Journal of Immunology, 193(2), 773–782. https://doi.org/10.4049/jimmunol.1400163

Zhao, X., Smartt, C. T., Li, J., & Christensen, B. M. (2001). Aedes aegypti peroxidase gene characterization and developmental expression. Insect Biochem Mol Biol, 31(4–5), 481–490. https://doi.org/S0965174800001557 [pii]

